# Integrated Open-Source Software for Multiscale Electrophysiology

**DOI:** 10.1101/584185

**Authors:** Konstantinos Nasiotis, Martin Cousineau, François Tadel, Adrien Peyrache, Richard M. Leahy, Christopher C. Pack, Sylvain Baillet

## Abstract

The methods for electrophysiology in neuroscience have evolved tremendously over the recent years with a growing emphasis on dense-array signal recordings. Such increased complexity and augmented wealth in the volume of data recorded, have not been accompanied by efforts to streamline and facilitate access to processing methods, which too are susceptible to grow in sophistication. Moreover, unsuccessful attempts to reproduce peer-reviewed publications indicate a problem of transparency in science. This growing problem could be tackled by unrestricted access to methods that promote research transparency and data sharing, ensuring the reproducibility of published results.

Here, we provide a free, extensive, open-source software that provides data-analysis, data-management and multi-modality integration solutions for invasive neurophysiology. Users can perform their entire analysis through a user-friendly environment without the need of programming skills, in a tractable (logged) way. This work contributes to open-science, analysis standardization, transparency and reproducibility in invasive neurophysiology.

## Introduction

Invasive electrode recordings are a unique source of in-vitro and in-vivo neurophysiological data at high resolution in both space and time, recorded in relation to complex animal and human behavior. The complexity of this kind of data has increased in recent years, with the advent of increasingly dense multi-channel and multi-site electrode arrays. This evolution provides exciting opportunities to explore the relationship between local events, such as action potentials, and more global dynamics at the systems level, such as fluctuations in oscillatory network activity. At the same time, these multiscale explorations require different analytical methods from those traditionally used in the field.

Challenges in exploring high-dimensional spatio-temporal data sets are not specific to electrophysiology: they occur frequently in neuroimaging data, as scanners produce increasingly large volumes of data, which are often shared across multiple groups or research centres. In response, the brain imaging community has made significant strides in developing shared software platforms to harmonize analytical methods and to facilitate data sharing (Abraham et al., 2014; Gorgolewski et al., 2011; Gramfort et al., 2013, 2014; Hanke et al., 2009; Tadel et al., 2011). Indeed, free, open-source software toolkits have been critical for facilitating training and augmenting research productivity. This approach has transferred to the field of scalp electrophysiology (Baillet et al., 2011), but as of yet it has not found widespread use in invasive neurophysiology (IN). Software tools do exist for specific segments of the IN data workflow, such as for spike detection and sorting and time-series analysis (Fee et al., 1996; Hazan et al., 2006; Hill et al., 2011; Mitra and Bokil, 2007; Oostenveld et al., 2011; Pachitariu et al., 2016; Quiroga et al., 2004), but they remain relatively specialized, some with limited support and documentation and most with restricted interoperability with other tools.

While we acknowledge significant efforts in harmonizing data formats for electrophysiology (Stead and Halford, 2016; Teeters et al., 2015; Neuroshare - http://neuroshare.sourceforge.net/index.shtml), it does seem that this field lags behind others in meeting the demands of recommended practices for data management and transparency (Gorgolewski and Poldrack, 2016; Larson and Moser, 2017). In this regard, well-supported software tools are required to produce analytical workflows that are validated, well documented and reproducible. Important components include data organization, review and quality control, verified implementations of signal extraction and decomposition methods, solutions for advanced visualization registered to anatomy, and sound approaches to machine learning and statistical inference. As in the brain imaging field, such tools would facilitate the reproducibility of published results and the dissemination of methods within and between research groups. They would also save considerable time and resources currently required to re-code published methods. In addition, re-coding presents challenges in code verification relative to a published method, raising possible concerns about the validity of the end results and limiting the long-term value of the effort.

Here we deploy and share open-source software (called *Invasive Neurophysiology*-Brainstorm, or *IN-Brainstorm*) that integrates multiple aspects of data analysis for most modalities and signal types for basic electrophysiology: from single cells to distributed channel arrays, from spiking events to local field potentials, from ongoing recordings to event-related responses, and from in vitro preparations to free-behaving models. We also emphasize the importance of an extensive graphical interface for user-friendly access to advanced analytical methods, of flexible scripting features for high-performance computing, and of traceable code execution. The proposed tool is accompanied by extensive online documentation and support from a user community web forum.

This free application builds on the foundations of the *Brainstorm* platform (Tadel et al., 2011), which is well-established (21,000 user accounts), free open-source software for magnetoencephalography (MEG) and electroencephalography (EEG). Brainstorm can integrate multimodal data volumes in addition to scalp electrophysiology e.g., magnetic resonance imaging (MRI), CT-scans and functional near-infrared spectroscopy (fNIRS). It also features advanced source modeling for electrophysiological signals.

The IN-Brainstorm application provides a comprehensive suite that interoperates with other, more specific and constantly evolving IN tools available from the open-source community e.g., for performing spike sorting. The end result is a unique and expansive software toolkit that bridges across recording scales and data modalities, registers invasive neurophysiology with structural anatomy data, and thereby delivers a unifying analytical environment to the neurophysiology research community.

## Results

The IN-Brainstorm functionalities described here offer comprehensive solutions for data importation and analysis, including spike-sorting, extraction of local field potentials, and correlations among these measures across multiple channels. Importantly, thanks to an intuitive graphical user interface, no programming skills are required for accessing and using the advanced methods available, including for assembling and sharing advanced data analysis pipelines. A summary of these software features is provided in Table 1, and a schematic of the workflow enabled by the toolbox is shown in FIGURE 1.

**Table 1.** Synopsis of the e-Phys toolbox and the tools that can be used for LFP analysis. The e-Phys toolbox provides a working frameworkfor every step of the e-Phys analysis and each module can easily be enriched with future additions.

| e-PHYS TOOLBOX |  |
| --- | --- |
| **Acquisition Systems** | Blackrock (.nsX), Ripple (.nsX), Plexon (.plx, .pl2), Intan (.rhd), Neurodata without borders (.nwb), Tucker Davis Technologies |
| **Spike Sorters** | WaveClus, UltraMegaSort2000, Kilosort (Klusters) |
| **LFP spectral Artifact Removal** | Bayesian spectral spiking-artifact removal |
| **Spike-related Functions** | Tuning Curves, Raster Plots, Spike Field Coherence, Spike triggered Average |
| LFP - ANALYSIS |  |
| **Pre-processing** | DC-offset removal, Band-pass / band-stop / notch filtering, resampling |
| **Artifact Removal** | SSP, ICA |
| **Frequency** | FFT, Welch Density, Morlet Wavelets, Hilbert transform, Multi-tapers, Phase Amplitude Coupling, Instantaneous frequency, Canolty Maps |
| **Connectivity** | Correlation, coherence, Bivariate Granger causality, Phase Locking Value, Amplitude Envelope Correlation, Phase Transfer Entropy |
| **Statistics and machine learning** | Parametric testing (zero / baseline), Partial Least Squares, Support Vector Machine classification, Linear Discriminant Analysis classification |

**Figure 1.**
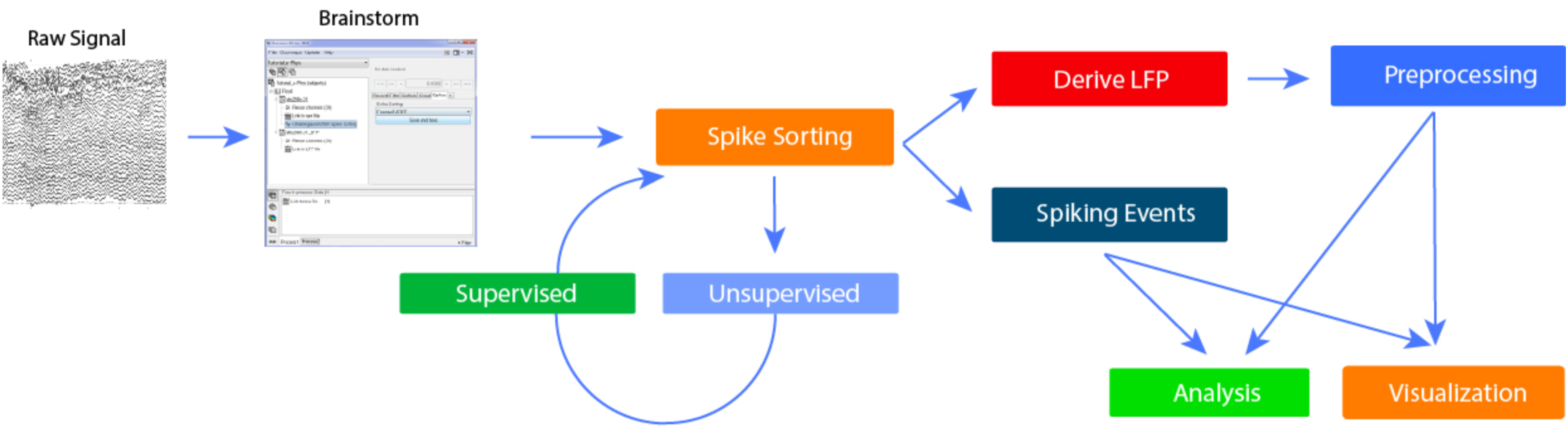
Workflow of the toolbox. Workflow of thee-Phys toolbox. Users initially import the header to the raw binary signal. Once the data is identified, the users perform the spike-sorting step. The spike-sorting process is divided into two parts: Unsupervised (the algorithm creates neuronal clusters automatically) and Supervised (the user inspects the output of the unsupervised part). At this point of the workflow, the LFP can be extracted. All the spiking events that were previously computed and the down-sampled LFP signals are all encapsulated to a single binary file. The original binary file can be stored to an external source and is no-longer needed. Finally, users can now perform preprocessing and analyze their data, utilizing the spike-related function that have been introduced to Brainstorm by this toolbox.

The bedrock of the present developments is the Brainstorm platform. Brainstorm (Tadel et al., 2011) is written in Matlab (Matlab2008a and higher) and Java. It is therefore independent of the operating system (Windows, MacOS and Linux). Community code management is via <u>GitHub</u>. Users without access to a Matlab license can use a fully executable version of the application compiled for the above operating systems. Extensive documentation is freely available online, with specialized tutorials, datasets and videos (https://neuroimage.usc.edu/brainstorm/e-phys/Introduction).

In the following sections, we describe a broad spectrum of analysis options for multiscale electrophysiology that are enabled by IN-Brainstorm and illustrate these features with the processing of an example raw data file.

### 1. Importing, reviewing and pre-processing raw data

#### 1.1. Raw data importation

Data to be analyzed must first be imported into the software. Brainstorm can read raw electrophysiology data from 80 different file formats. We have added new data formats specific to single- and multi-unit electrophysiology, including Plexon (.plx and .pl2), Blackrock (.nsX), Ripple (.nsX), Intan (.rhd), Tucker Davis Technologies, and Neurodata Without Borders (.nwb). New formats can be added on demand. Raw data can also be read directly from ASCII and basic binary data formats, with header file parameters easily specified from a GUI.

#### 1.2. Data review

Raw files of continuous data from chronic preparations can be voluminous due to hours-long durations, tens of kilo-Hertz sampling rate and simultaneous recording from multi-channel electrode arrays. Hence loading such large raw files at once into computer memory can be impractical. For this reason, we have implemented efficient data review solutions that load portions of the data on the fly, depending on the visualization parameters set by the user (e.g., virtual page length, selection of a subset of channels or montages for review, keyboard and mouse shortcuts for navigating and marking events).

Task events (e.g., stimulus types and presentation times, behavioral responses) and ancillary recordings (electrooculograms, electrocardiogram, eye and body movements, video recordings of behaviour, etc.) are readily registered to the electrophysiological data in IN-Brainstorm, for multimodal data review, quality control and event-related processing. We emphasize that when a raw file is reviewed, the physical data is not duplicated as a Brainstorm file. Instead, the header of the original data file is automatically parsed to extract metadata, such as channel parameters, sampling rate, time stamps, event codes, etc.

Figure 2 (left) shows an example of IN-Brainstorm display for data review, including sub-menus for displaying and navigating through files and events. The right panel shows an example of raw data collected with a Plexon MAP system and a 32-channel linear electrode implanted in cortical areas MT and MST of a non-human primate. The animal maintained fixation during the presentation of a motion stimulus comprising of dots that translated in 8 different directions.

**Figure 2.**
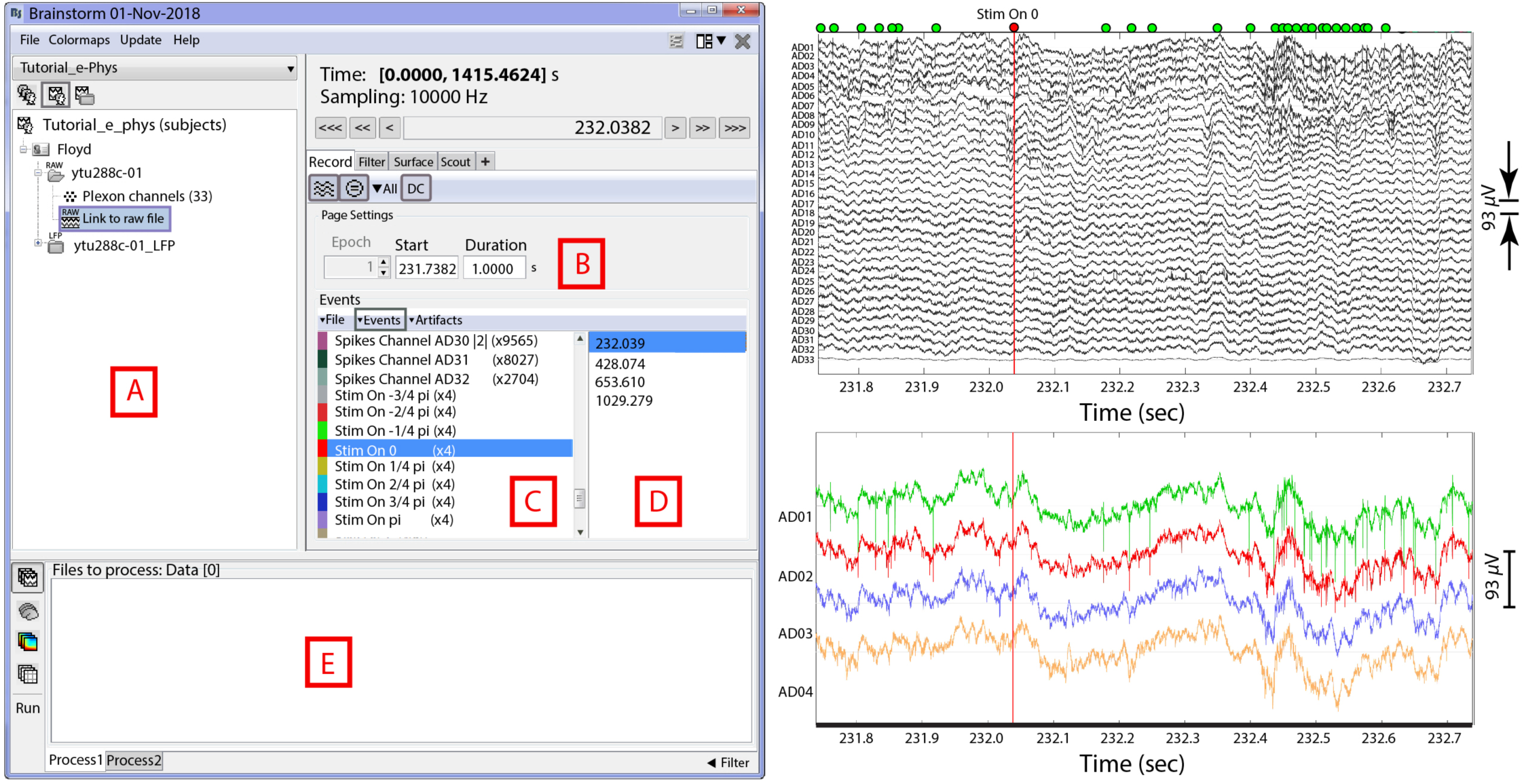
Dataset navigation and pre-processing window. Left: (A) Main Brainstorm window showing the created dataset entry (ytu288c-01) on the data-tree section. (B) Selection of starting time point and duration ofsignal segment to be loadedfor reviewing. (C) Experimental and spiking events are displayed. (D) An event is selectedfrom the “Stirn on 0” condition. This selection automatically synchronizes all reviewing windows to the timepoint of the event’s occurrence. Right Top: 1 second segment displaying raw signals from all electrodes. The vertical red line indicates alignment around the selected event (“Stirn On 0”). The green dots on the top of the figure represent the spiking events from the first neuron on electrode with label AD01. Right Bottom: A selection of the first 4 electrodes, aligned in time with the top figure.

The red line in the figure shows the time of a *Stim On* event, extracted from the data. Spikes detected online (labelled as *Spikes Channel*) were extracted directly from the raw file contents by IN-Brainstorm, with automatic registration to the data time series.

The bottom right panel of Figure 2 shows a selection of 4 channels temporally aligned with the top figure. The spikes from a neuron that was isolated on the first electrode are marked with green circles at the top of the full time-series displayed in the top panel. Users can browse the raw traces using point-and-click GUI and a series of keyboard shortcuts. On-the-fly bandpass and notch filtering can be applied to the signals.

#### 1.3. Quality control & data pre-processing

Starting from the kind of raw data shown in Figure 2, users can easily navigate through the recordings and experimental trials and events for quality control. Data segments, channels and entire trials can be marked as “bad” and excluded from further analyses using automatic processes or based on user evaluations.

The IN-Brainstorm pre-processing toolkit features solutions for adjustments of recording baseline, data resampling and frequency filtering (with linear phase filters). Additionally, detection and attenuation of artifacts (e.g., heartbeats, eye and body movements, stimulation and juice artifacts) can be achieved with principal (Uusitalo and Ilmoniemi, 1997) or independent component analysis (Bell and Sejnowski, 1995; Cardoso, 1999). Finally, combining sensor data with the actual geometry of the recording array(s) enables many 2-D and 3-D visualization possibilities for time-series and realistic topographical plots, as illustrated further below.

### 2. Spike detection and spike sorting

Following the importation and preprocessing of data, IN data is often processed to extract spiking events from single or multiple neurons. This entails detecting spike occurrences and classifying these events according to their respective neural sources (Quiroga, 2007). Most data acquisition systems feature online spike detection and sorting. These online events can be imported directly into IN-Brainstorm with the corresponding raw recordings. Yet, usual IN practice is to refine spike classification with a two-step procedure consisting of 1) unsupervised clustering, which automatically assigns each spike to a neural source based on waveform features, then 2) supervised clustering, which requires manual reviewing and editing of the labels from unsupervised clustering and the elimination of spurious spike events.

For IN-Brainstorm, we have enabled the direct interoperability with a selection of existing and openly-available spike-sorting toolkits: *Waveclus* (Quiroga et al., 2004), *UltraMegaSort2000* (Fee et al., 1996; Hill et al., 2011) and *Kilosort* (Pachitariu et al., 2016). Those packages can be downloaded and installed automatically, in a completely transparent procedure. Sequentially, these tools are called by and interact with IN-Brainstorm without programming interventions from users.

#### 2.1. Unsupervised spike sorting

Figure 3 (left) shows IN-Brainstorms’ GUI for unsupervised spike-sorting. Raw files are dragged and dropped into the GUI process box before a spike-sorting tool is selected from the IN-Brainstorm toolkit. Next, spike events are detected on each electrode and classified according to their putative neuronal generators.

**Figure 3.**
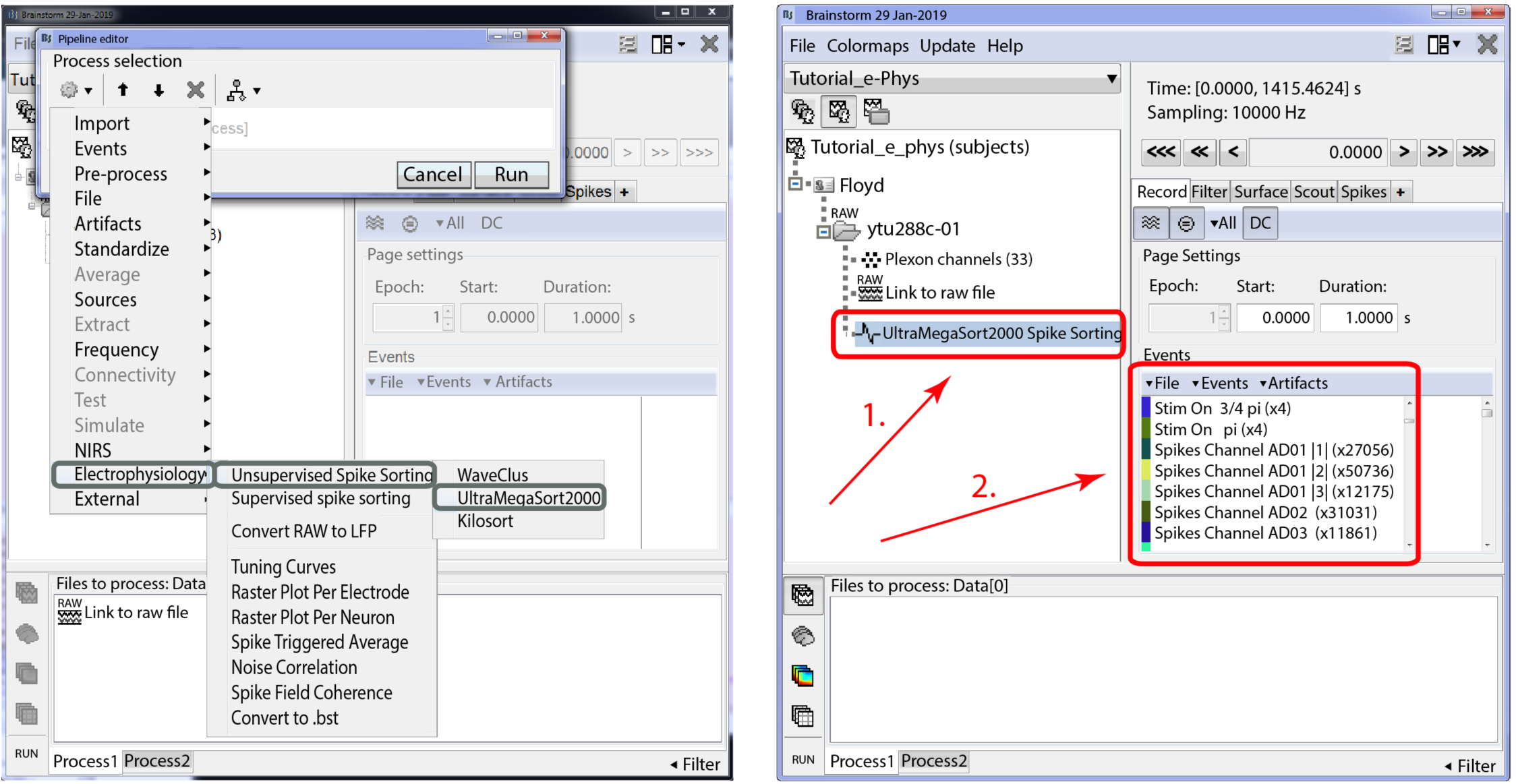
Unsupervised Spike-sorting. Left: Selection of the embedded spike-sorters for unsupervised spike-sorting within Brainstorm. Right: Example of a dataset spike-sorted with UltraMegaSort2000. (Box 1) A new entry in the database UltraMegaSort2000 Spike Sorting appears and indicates that this dataset has been spike-sorted. (Box 2). New events appear in the Events window, corresponding to the spikes that the spike-sorter clustered (Spikes Channel X).

The unsupervised spike events produced overwrite the online counterparts that were detected during data acquisition. The output of the spike-sorting process (Figure 3 Box 1) is automatically registered to and accessible from the IN-Brainstorm database and linked to the corresponding raw file. The spike events are labelled in a principled manner (per channel and source cell number – Figure 3 Box 2).

#### 2.2. Supervised spike sorting

As *WaveClus* and *UltraMegaSort2000* have built-in supervised spike sorting graphical user interfaces, we synchronized their GUIs with IN-Brainstorm’s. For *Kilosort*, we developed specific GUI bridges via *Klusters* (Hazan et al., 2006). The user-selected supervised clustering tool is called from Brainstorm’s main window after an unsupervised spike-sorted file is selected (Figure 4A). The user then switches to the GUI of the selected supervised spike clustering tool (Figure 4 B-D). Once supervised spike clustering is complete, the spike events are updated accordingly and registered into the software’s file system. Double-clicking on the link to the raw data file lets the user review the updated spike events along with the raw electrophysiological traces as shown in Figure 2 (Right).

**Figure 4.**
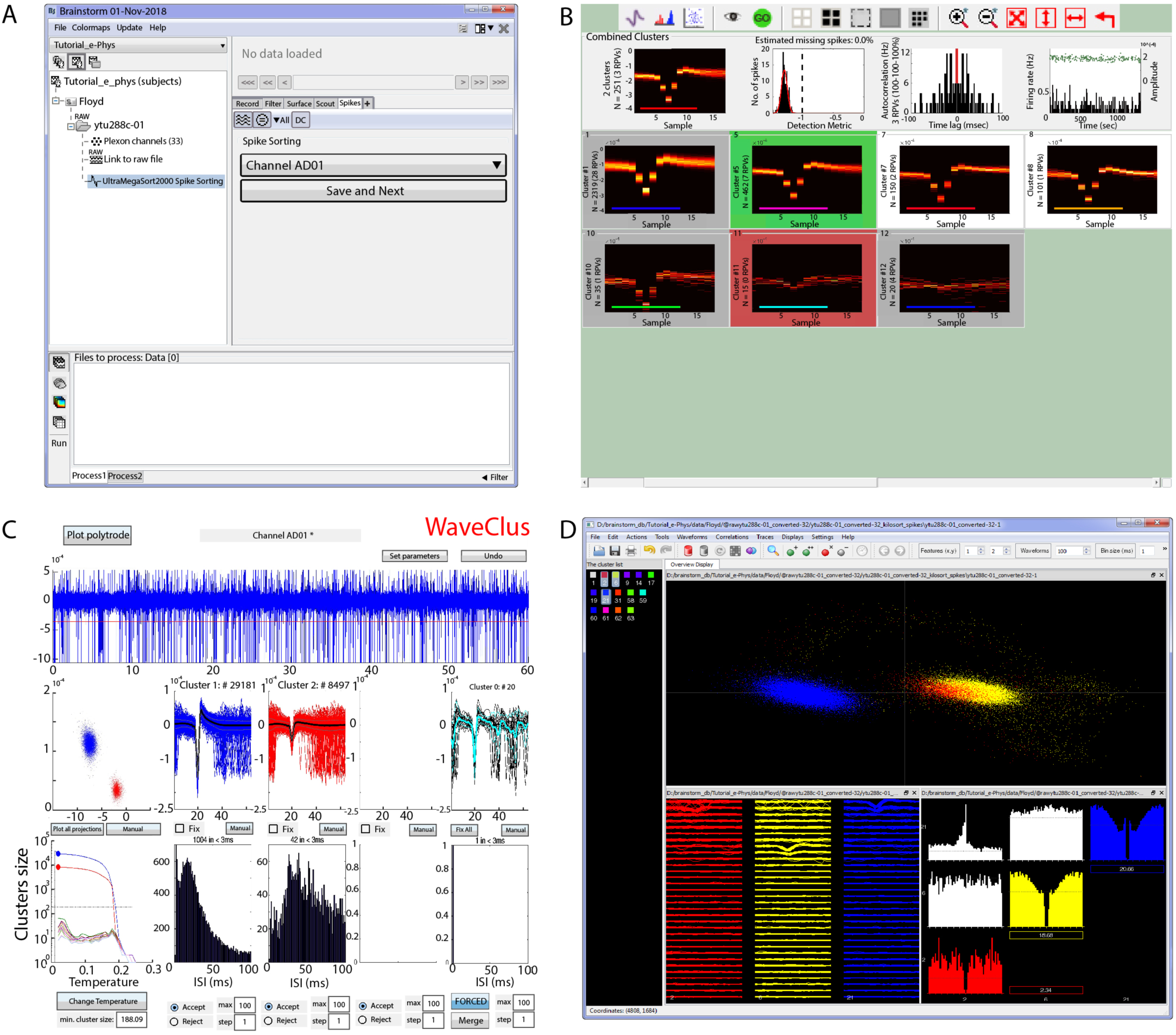
Supervised spike-sorting. (A). Main window that the user selects the electrode (or group of electrodes for Klusters) and the spike sortedfiles automatically update the spike sorter in use. Once the neuronal clusters have been adjusted, the user presses the Save and Next button and the next electrode gets selected to continue the supervised spike-sorting. (B, C, D). Supervised GUI for UltraMegaSort, WaveClus and Klusters respectively.

Spike events and categories from other spike-sorting tools can be readily imported as Brainstorm events, following the procedure described in the online documentation (https://neuroimage.usc.edu/brainstorm/e-phys/ConvertToBrainstormEvents).

### 3. Extraction of local field potentials

In addition to spiking activity, IN recordings yield local field potentials (LFPs), which provide direct measures of the summed post-synaptic electrical activity in the vicinity of recording electrodes (Legatt et al., 1980). These can be useful as a complement to spiking activity or a surrogate for some aspects of neural activity (e.g.,(Mineault et al., 2013)), provided that LFP traces can reliably be filtered and separated from spike waveforms (Zanos et al., 2011).

Figure 5A shows the IN-Brainstorm’s GUI for extracting LFP traces from raw recordings. The application features efficient tools to remove spike traces (Zanos et al., 2011), to perform anti-aliasing bandpass filtering and to down-sample the raw data. The de-spiking method proposed by Zanos et al. (2011) increases the accuracy of subsequent spike-field coherence measures and of spike-triggered average signals.

**Figure 5.**
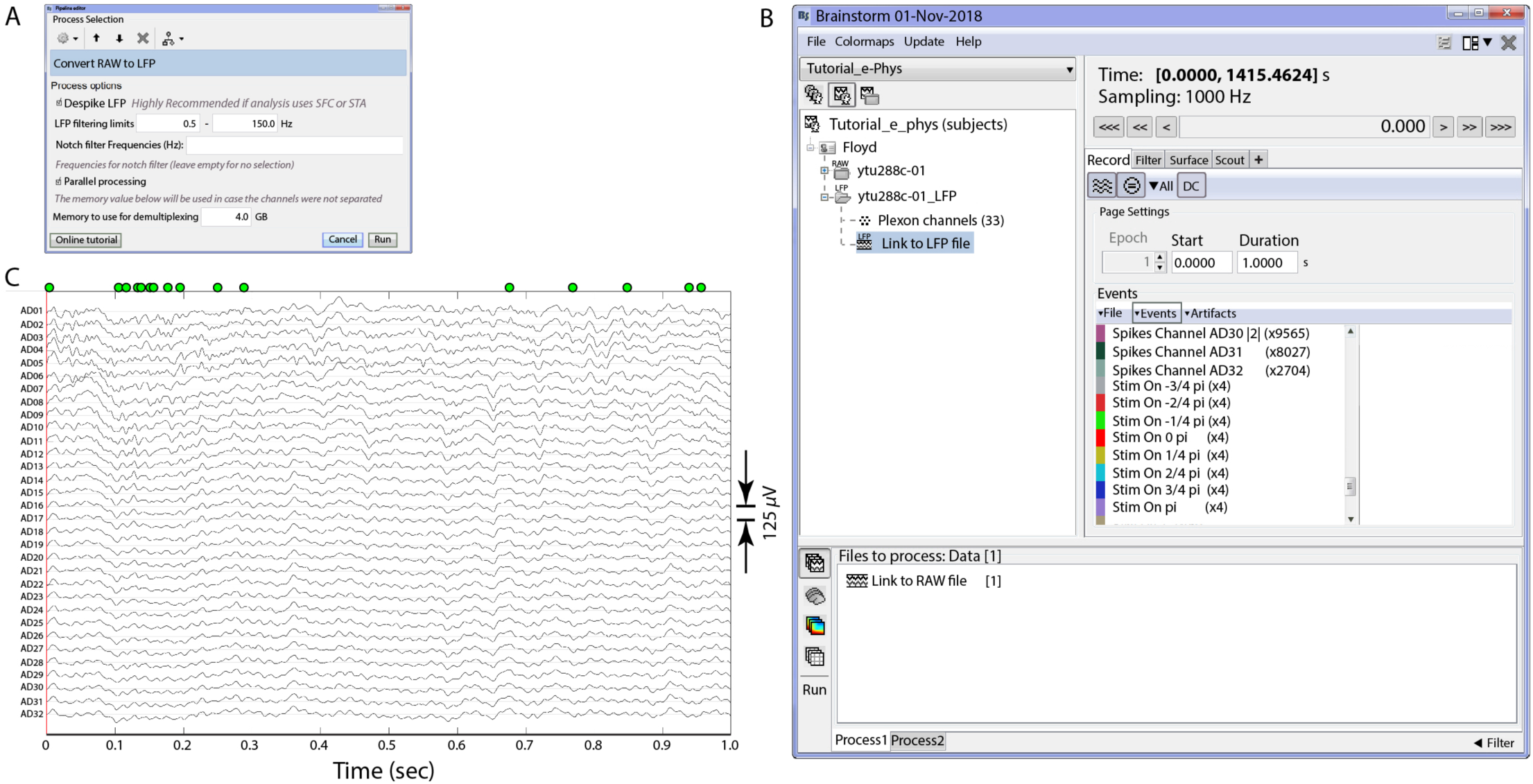
Conversion from raw signals to LFPs. Converter from raw signals to LFPs (A). Users can select the filtering limits of their LFP and apply a notch filter if necessary. The converter downsamples the raw signal to 1 KHz. (B). Once the conversion is complete, a single binary file (.bst) that contains all the necessary information (LFPs, experimental and spiking events) is stored on the hard drive, and automatically imported on the data-tree as a new dataset. (C). 1 second segment review of the created LFP signal traces. The spiking events from the first neuron of electrode AD01 are represented by the green dots on the top of the figure, as in Figure 2.

The resulting LFP traces and experimental events are automatically registered in IN-Brainstorm’s data repository for further review and analysis with a vast library of tools and pipelines - as described below - or for easy exportation to other software or plain files.

LFP extraction produces a new IN-Brainstorm down-sampled time-series binary file (Figure 5B) with all the corresponding metadata, such as channel description (e.g., electrode labels and locations), and spike and experimental events. This file is easily sharable among researchers since its size is typically ∼20-30 times smaller than the original raw file. Figure 5C shows a segment of the LFP file created.

### 4. Epoching

Once the relevant neural signals (LFPs and spikes) have been extracted from the raw data, they can be divided according to experimental epochs. Epochs are typically comprised of experimental trials, with the time window selection defined around a stimulation or behavioral event of interest. These can be imported directly into the IN-Brainstorm file system.

To illustrate these functions, we make use of the example visual cortex recording described previously (Figure 2). The experiment involved presentations of moving stimuli while the animal maintained fixation; we defined the relevant epochs as segments of [-500, 1000] ms around the onset of each visual stimulus (Figure 6 Left). In total we considered 8 different directions of the visual stimulus moving pattern; each stimulus condition was repeated 4 times (one condition was repeated for 96 trials for usage in the raster plot, and noise correlation functions). Imported trials to the database are shown in (Figure 6 – Right).

**Figure 6.**
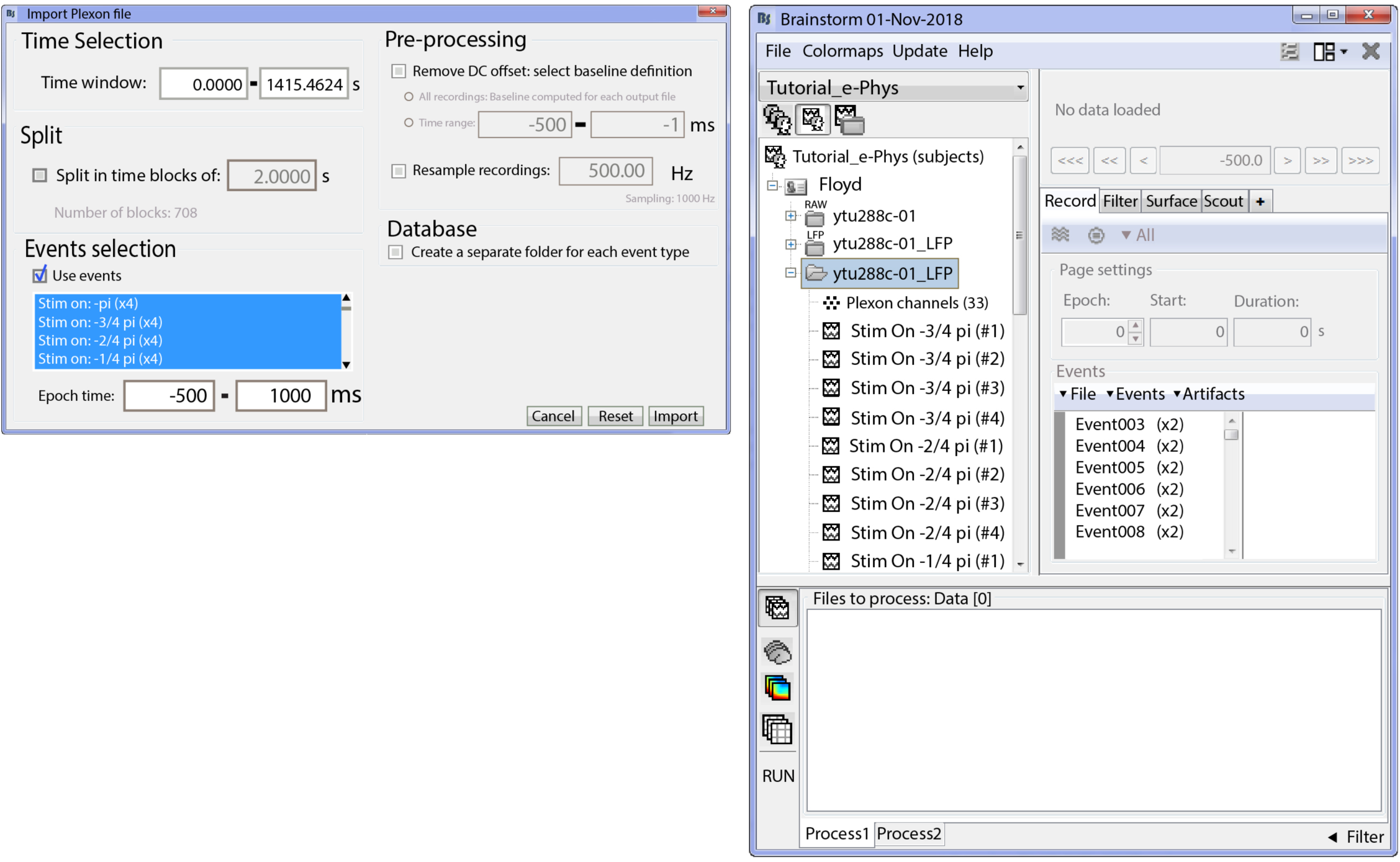
Importing of trials. (Left) Selection of the events of interest and temporal boundaries around them, for importing the LFP segments. (Right) Imported LFP trials for the selected conditions in the Brainstorm database.

The following analysis steps can then be applied on the epoched trials.

### 5. Analysis of individual LFP signals

LFP traces can be analyzed using Brainstorm’s extensive library originally developed for EEG and MEG research (Tadel et al., 2011). We show in Table 1 a list of the main data processing categories that are available for LFP analysis. There is extensive online documentation, accompanied by data files, that describes in detail the methods and practices of LFP signal analysis (http://neuroimage.usc.edu/brainstorm).

We briefly provide below a few examples of these functions and their implementation in IN-Brainstorm.

#### 5.1. Time-frequency decompositions

Having extracted the LFP signal and defined an appropriate analysis epoch, one can compute the LFP power at different frequencies and at different times relative to a stimulus event. Such information is often used to infer stimulus selectivity, anatomical sources of input, and other factors that are not necessarily apparent in spiking activity (Buzsáki, 2006; Fries et al., 2008; Pesaran et al., 2002; Wilke et al., 2006; Womelsdorf et al., 2006).

IN-Brainstorm provides functionality for spectral and time-frequency decompositions, which can be derived using power spectrum density estimates, Hilbert or wavelet transforms. An example time-frequency decomposition (wavelet) is shown in Figure 7A for the example LFP data corresponding to a single stimulus condition and epoch that shows a strong beta response after stimulation. The wavelet decomposition was z-scored with respect to a pre-stimulus baseline [-500:-100] ms.

**Figure 7.**
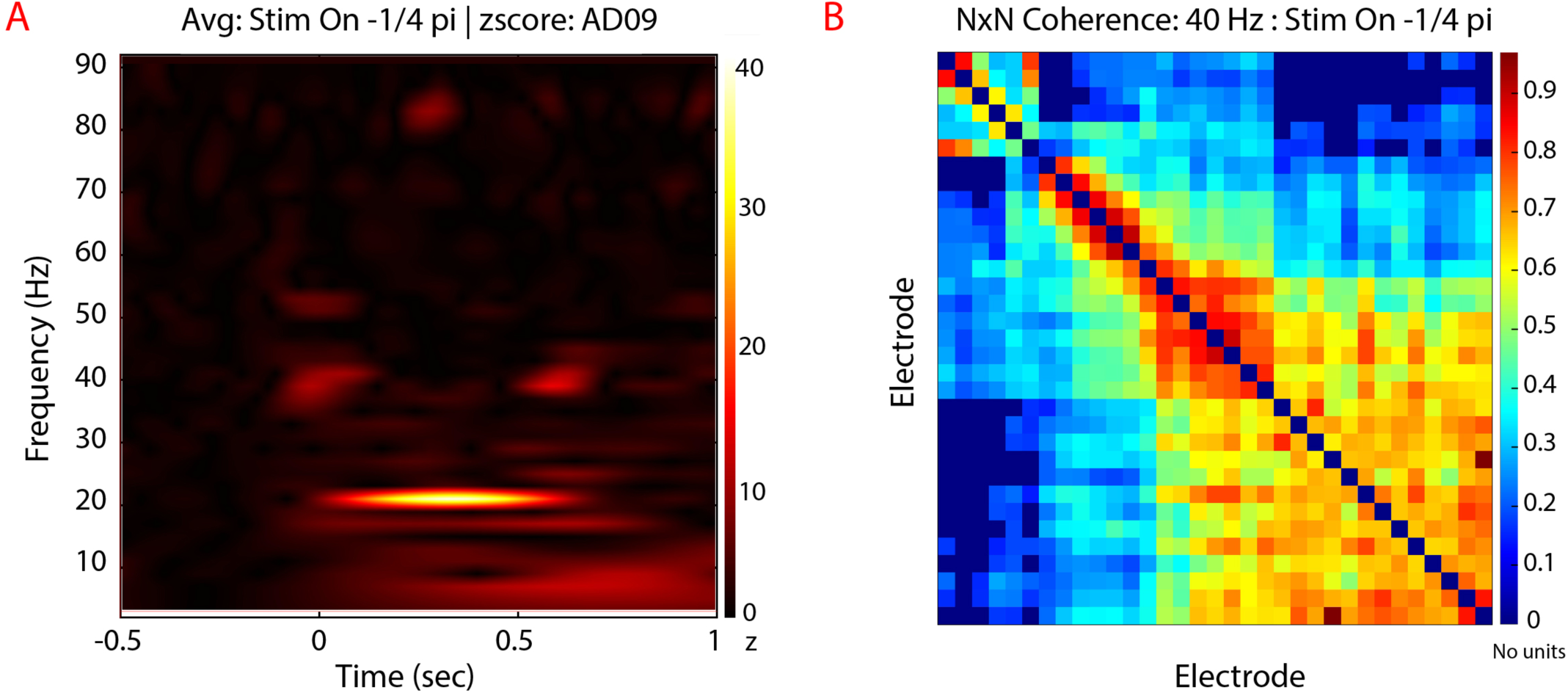
Analysis of LFP signals. (A). Wavelet Decomposition ([2-90] Hz) of a single LFP trialfrom electrode labeled AD09 for moving stimulus condition towards −1/4 pi degrees direction. Users can select the channel they want to be displayedfrom a drop-down list on the main Brainstorm window. (B). Estimation of NxN coherence for a single trial across all electrodes. The coherence values are co/or-coded for a specific frequency (on the example 40 Hz frequency is selected). Users can display coherence in the otherfrequencies by moving a sliding toggle.

#### 5.2. LFP-LFP signal analysis

LFP signals from multichannel recordings can be analyzed to detect occurrences of various forms of signal similarities in the time or frequency domain. These measures are often interpreted as representing functional connectivity between different sites (Fries, 2005; Fries et al., 2002, 2008; Womelsdorf et al., 2006). IN-Brainstorm provides support for widely-used measures based on amplitude or phase statistics as indicators of possible interregional brain interactions (coherence, phase-locking values, bandlimited amplitude envelope correlations, phase-transfer entropy) and parametric models (estimates of time- or frequency-domain Granger causality). Advanced measures of interdependence between oscillatory components of polyrhythmic brain activity can be derived with phase-amplitude coupling (PAC) estimation tools (Canolty et al., 2006, Samiee and Baillet, 2017). An example estimation of coherence among all combinations of electrodes is shown in Figure 7B for a single stimulus condition and epoch. The bimodal pattern that emerges (high coherence among some channels and low coherence among others) is an indication of the transition of the linear probe across neighboring cortical areas, from MT (electrodes 1:13) to MST (14:32).

### 6. Analysis of individual neuron spiking activity

Spikes are registered in IN-Brainstorm as events; the corresponding features are 1) the time of occurrence and 2) a label for distinguishing between neuronal sources. We provide several features for visualization of epoched spiking data.

#### 6.1. Raster plots

Raster plots are routinely used to visualize the relations between neuronal firing and a stimulus event or a behavioral response. The spike occurrences in each trial and from each neuron are binned over user-defined time epochs and converted into firing rates (spikes/second).

We provide two methods for visualizing spiking activity with IN-Brainstorm:

The first method shows the spiking data as trial vs. time bins for each neuron. Raster plots of spiking rates are displayed after interactive selection of the cell to be reviewed. Figure 8A shows the raster plot of the first neuron detected from contact AD01, with 10-ms binning. The trial average of the neuron’s firing rate from 96 trials of a single condition revealed a stimulus-onset-to-maximum-firing latency of about 150 ms.

**Figure 8.**
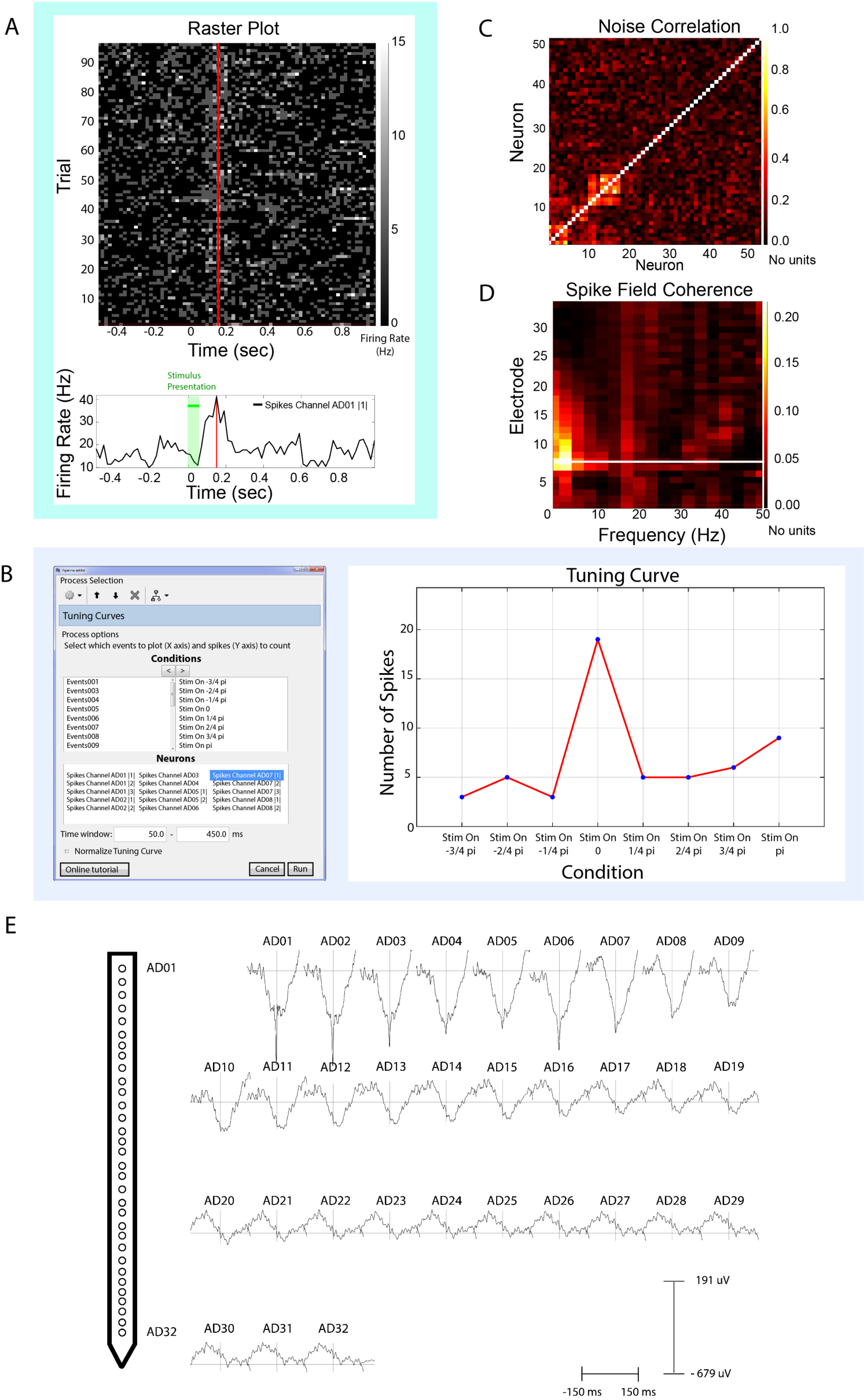
Spike-LFP analysis functions. (A) Raster Plot. (Top) Example raster plot for 96 trials of a single condition for the first neuron picked up on electrode labeled ‘AO0l’. Firing has been binned into lOms segments. The green shading indicates the period where the stimulus was presented on the screen. A single condition was repeated 96 times instead of 4 for the purpose of this raster plot. (Bottom) Averagefiring rate from all trials shows maximum firing ∼150ms after the stimulus onset for this neuron. The vertical red line indicates interactive temporal alignment between the two plots and the green shade the period of the stimulus presentation. (B) Tuning curves function. (LEFT): Users select the neurons that they want to display the tuning curves, and sequentially the conditions (and their order) that would formulate the x-axis (right side of the conditions selection window). Additionally, we included a selection for the time-window where the spikes would be counted. (RIGHT): Tuning curve for an example neuron, selectedfrom the window on the left of Figure B. The x-axis shows the different experimental conditions at the order selected on the previous window. This neuron expresses selectivity for the condition “Stirn On 0”. (C) Noise correlation. The function selects all the neurons that elicited spikes within the trials imported and displays a nxn figure where the noise correlation is computed for all combinations of neurons. Specifically for the dataset illustrated, there were 53 unique neurons picked up by the electrodes (according to the spike sorting step). This figure shows the computation of noise-correlation on all trials (a subset on a specific condition is also possible) for the presentation 96 trials of a motion stimulus, and spikes are selected on [0,300] ms around the stimuli presentations. (D) Spike field coherence for an example neuron picked up from the first electrode (AD07) for the motion stimulus condition “Stirn On 0”. The spike-field coherence window displays spectral influence of a single neuron to all 32 electrodes. Frequency is shown up to 60 Hz. Time selection around each spike was [-150, 150] ms (E) Spike triggered average of a neuron picked up on electrode labeled AD0l. A graph of the linear probe with the relative electrode locations is displayed on the left of the figure. The time selection around the spikes was set to [-150,150] ms for all trials of all experimental conditions. The electrodes have been presented into four rows for easier visualization. In reality, the linear probe extends on a single dimension (1:32). The scale of the STA is shown on the bottom right. All traces have been aligned to the same time-selection (0 ms - time occurrence of the spikes of AD0l}, indicated by the vertical line of each signal’s display.

The second method is embedded within the topographical plots section as shown below.

#### 6.2. Tuning curves

Tuning curves capture the relationship between an experimental variable (e.g., the orientation of a visual stimulus) and a scalar measure of neural activity (e.g., a single neuron’s trial-averaged firing rate).

Tuning curves are readily produced from continuous data files that contain the event markers of interest to the study. Tuning curves are displayed with IN-Brainstorm after manual assignment of the order of the experimental conditions (x-axis), the selection of the neurons to be displayed, and the selection of the time window of interest for reporting spiking activity. A separate tuning curve figure is produced for each neuron selected.

We selected the events and individual neurons previously identified from spike sorting via IN-Brainstorm’s GUI. Figure 8B shows the tuning curves of one example neuron (labeled as “*Spikes Channel AD07 |1|”*) for the 8 different conditions (Stim On −3/4 pi, Stim On −2/4 pi etc.) of the motion stimuli. The tuning curve shows the preference of this neuron for stimuli moving in the right direction (*Stim On 0* condition).

#### 6.3. Topographical plots

When multichannel recording devices are used, neurophysiology data can be shown as topographically registered to structural anatomy. IN-Brainstorm can show neuronal firing at the 3-D locations of the recording probes/arrays. To illustrate this feature, we used a separate dataset that was collected from two 96-channel Utah arrays and one 32-linear probe (Krause et al., 2017). A structural T1-weighted MRI volume was acquired preoperatively. The head and brain surface envelopes were segmented with Freesurfer (Fischl et al., 2001) and directly imported in IN-Brainstorm. The electrode contact locations were co-registered to the 3-D anatomical volume by specifying the distance of the electrodes along the probe and locating the tip of the probe and the entry point through the skull, using Brainstorm’s MRI volume viewer.

Neuronal firing was binned in 10-ms segments and displayed on the animal’s anatomy as shown in Figure 9A (a single bin is displayed in the figure). This figure shows IN-Brainstorm’s ability to overlay the segmented cortical surface, MRI orthogonal slices, the implanted devices with actual geometry, and color-coded displays of raw or processed electrophysiology data (here instantaneous firing rates). Figure 9B shows a zoomed-in version of Figure 9A over the Utah array implanted in the prefrontal cortex.

**Figure 9.**
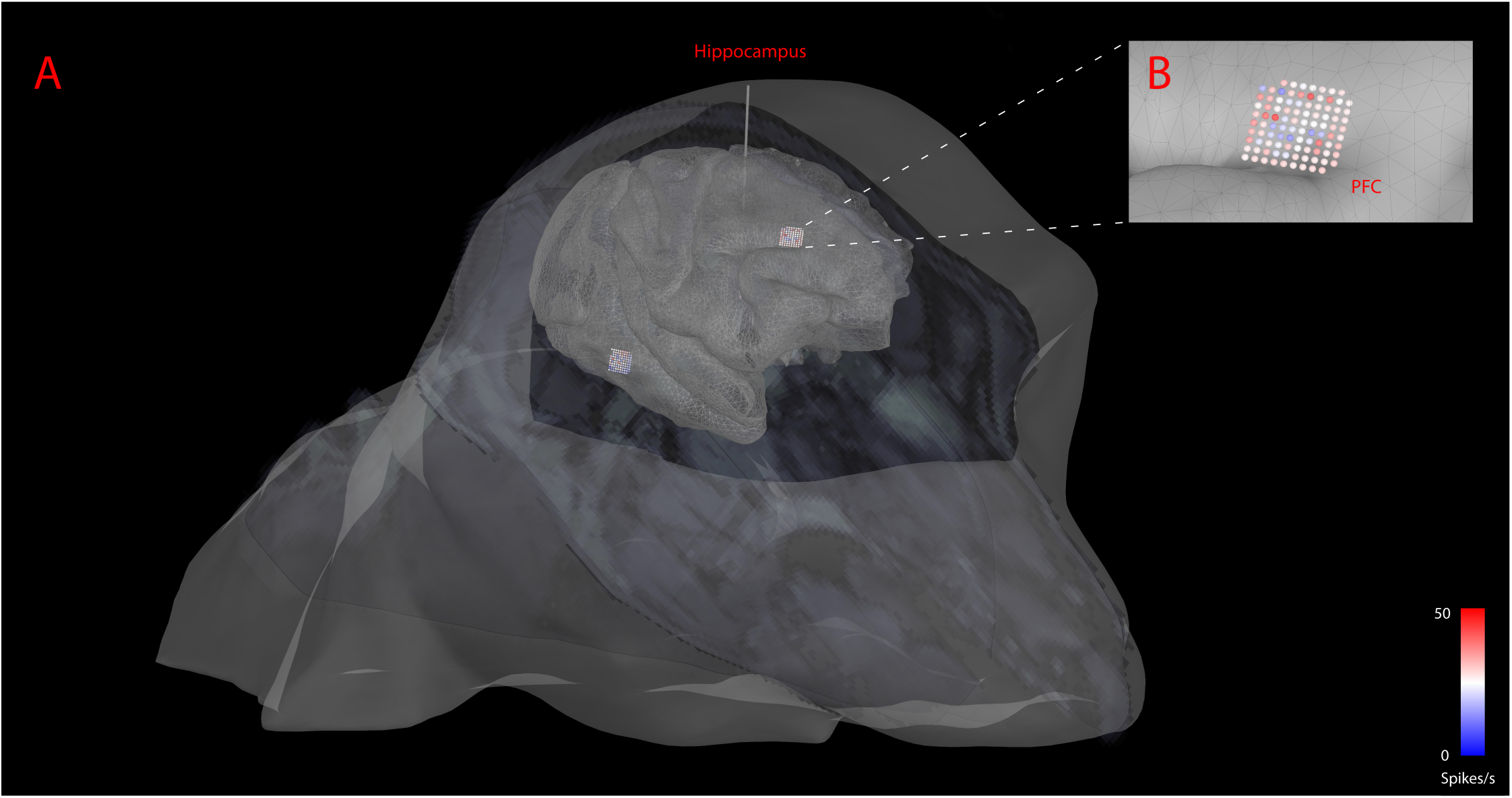
Topographical visualizations. (A). Visualization of the implanted electrodes on the anatomical MRI of the non-human primate. The MR slices are superimposed on the figure. (B). Zoomed in version of an implanted Utah array on the cortical surface, with the spiking activity color-coded on the array’s topography for a single time-bin.

### 7. Spike-spike analysis: Noise correlations

While tuning curves capture neuronal sensitivity to stimulus properties, the fidelity of a population code is thought to be limited by noise that is common across neurons (Zohary et al., 1994); for example, neurons would be noise correlated if for each stimulus their activities are correlated (Eyherabide and Samengo, 2013). Such noise correlations are typically quantified as the Pearson correlation coefficient between the firing rates of two neurons across trials. Such correlations strongly influence the accuracy of population coding (Abbott and Dayan, 1999; Averbeck et al., 2006; Liu et al., 2016; Panzeri et al., 1999; Sompolinsky et al., 2001).

Noise correlation statistics are displayed with IN-Brainstorm from the correlation of the spike trains that each neuron elicited within a given epoch, for all neuronal combinations. The end result is a *n*x*n* matrix (with *n* the number of unique neurons that produced spikes during the selected trials) that shows noise correlation estimates between the selected neurons.

Figure 8C shows the noise correlation profile across the 32-channel array of the example dataset, for 53 unique neurons that elicited spikes across all trials at the 8 conditions of presentation of the moving stimulus in the original data set from Figure 2. Spikes included in the correlation computations were selected in the [0,300]-ms time range of each trial.

The computed noise correlation showed 2 pairs of neurons with abnormally high noise correlation (above 0.8). After further inspection, it was revealed that this was due to the fact that the spike-sorter that was used was not taking into account the relative position of the electrodes, and the same neurons were picked up from neighbouring channels:

Neurons: AD01 |1| - AD02|2| and AD08 |1| - AD09 |1| were the same neuron.

### 8. Spike-LFP analysis

Spikes are local events, reflecting outputs from individual neurons. LFPs in contrast can capture activity over regions, including subthreshold post-synaptic activity, and therefore reflect the state of a broader network (Cui et al., 2016). There is considerable interest in relating the two types of signals for estimating the dependence of spiking activity on the broader context in which the neuron is embedded.

#### 8.1. Spike-field coherence

Spike-field coherence (SFC) estimates the consistency between the time occurrence of spike trains and the phase of co-localized LFP cycles as a function of frequency (Arce-McShane et al., 2018). SFC can also be used to evaluate synchronized activity between distant brain regions, as a marker of neuronal communication (Fries, 2005; Gregoriou et al., 2009; Liebe et al., 2012; Singer, 1999; Womelsdorf et al., 2007). IN-Brainstorm features the spike-field coherence estimator proposed by Fries et al. (2001). The user can derive SFC estimates for each GUI-selected neuron, for all electrodes and frequencies of interest.

Figure 8D shows SFC up to 60 Hz between a single neuron detected at channel AD07 of the example data set and the LFP traces at all the 32 channels of the probe. The time window selected around the spiking events was [-150, 150] ms. The horizontal white line indicates the electrode where the neuron was detected.

#### 8.2. Spike-triggered average of the LFP

Spike-triggered averaging (STA) of the LFP reveals how neuronal spiking is related to the dynamics of proximal or distant LFPs (Jin et al., 2008; Nauhaus et al., 2009; Ray and Maunsell, 2011). STA proceeds with trial averaging of LFP traces time-locked to a designated neuron’s spike events, followed by normalization with the total spike count.

Analogous to spike-field coherence, STA is computed over a user-selected time window around each spiking event. STA scores are per neuron, showcasing the average LFP amplitude around the occurrence of the spikes of each neuron. STA can be visualized on topological 2-D representations of the recording array, to reveal time-locked associations between neuronal spiking activity and local or remote LFP recordings.

Figure 8E shows the STA time-locked to the firing of the first neuron detected by electrode AD01 across trials and conditions. The topographical 2-D plot is produced with IN-Brainstorm using multidimensional scaling of the actual 3-D location and geometry of the implanted probe. The LFP epoch around spike event was [-150,150] ms.

### 9. Statistical inference and machine learning

Once measures have been extracted from spiking or LFP data, tools to conduct inferential statistical analysis in the multiple dimensions of electrophysiological data (space, time, frequency, connectivity) are available from Brainstorm’s library.

Parametric (one- and two-sample tests) and nonparametric permutation tests, descriptive and distribution statistics from histograms (Q-Q plot and Shapiro-Wilk test for data normality) are available. Here too, the software architecture emphasises interoperability with other toolkits, for expanded resources. For instance, multidimensional and nonparametric cluster statistics can be run on LFP and time-frequency data, from Brainstorm, via calls to FieldTrip (Oostenveld et al., 2011).

In addition, statistical learning tools for decoding and multivariate pattern analysis (MVPA) are also available (see e.g., Cichy et al., 2014). The Brainstorm library also includes support vector machine (SVM) and linear discriminant analysis (LDA) classification of LFP time series based on experimental events and conditions.

### 10. Additional features

#### 10.1. Processing power

Hardware acceleration in the processing of long recordings is enabled by Matlab’s standard parallel computing (e.g., multi-core) features, which are controlled directly from Brainstorm’s GUI. Flexible management of memory resources is also accessible to users, with the specification of the amount of RAM allocated to data manipulations while executing the LFP extraction process.

#### 10.2. Data management

Generally speaking, formal data management plans are seldom adopted by electrophysiology labs. Instead, the handling of data is typically project-based, with trainees managing their individual data collection and analyses until publication. When they move on to another project or to the next step of their career, they frequently leave data, analysis pipelines and results behind, with minimal documented organization for sustainability and knowledge transfer. This limits the long-term value of data and negatively impacts the reproducibility and verification of research results (Baker, 2016). Brainstorm has tools to improve and facilitate data management: data is hierarchically organized by Studies, followed by Subjects/Samples and (experimental) Conditions, which point to data elements such as links to raw data files, single-trial epochs, sample statistics, and other derivatives: power spectra, wavelet decompositions, measures of cross-frequency coupling and inter-regional connectivity, etc. As with all features in the application, user interactions with Brainstorm’s data organization are facilitated both by the application’s GUI and direct access via scriptable functions using Matlab code.

Another important aspect of Brainstorm is its capacity for importing entire data repositories at once, with associated metadata, when those datasets are organized according to the emergent Brain Imaging Data Structure (BIDS). Originally driven by the neuroimaging community, BIDS is a grassroots effort to harmonize data organization and documentation (Gorgolewski et al., 2016). BIDS has recently been extended to MEG electrophysiology (Niso et al., 2018) and is presently integrating EEG (Pernet et al.), and invasive neurophysiology (Holdgraf et al.).

#### 10.3. Batch processing

The software has a specific GUI for assembling data processing pipelines in an intuitive manner, choosing elementary processes from the (IN-)Brainstorm library and assembling them together into a logical progression along the workflow. These pipelines enable the reproduction of any data workflow with a click of a button. They can also be shared in Matlab format with collaborators or the entire user community. The Matlab code for pipelines can also be generated automatically by Brainstorm e.g., for execution in headless (no GUI) mode on high-performance computing servers and cloud resources.

## Discussion

We provide a free, extensive open-source software application for invasive electrophysiology. IN-Brainstorm is built on the foundations of Brainstorm, which was originally designed for human multimodal electrophysiology and imaging. IN-Brainstorm supports multiple data formats of raw signals from a variety of acquisition systems. The recorded traces and their LFP versions can be reviewed, quality-controlled and processed within a unique analytical environment, with easy GUI interactions, rich visualization, intuitive pipeline editing for scripting and sharing. We have built bridges for IN-Brainstorm to interoperate seamlessly with established, free spike-sorting tools.

A specific emphasis was put on providing versatile solutions for multidimensional data visualization, including 2-D and 3-D topographical plots registered to structural anatomy from co-registered MRI data. Source modeling of array data is also available using boundary element modeling of head and brain tissues (Gramfort et al., 2010; Kybic et al., 2005) and a variety of source modeling techniques available in Brainstorm (Baillet et al., 2001). Videos synchronised to electrophysiological traces can also be imported and visualized simultaneously in synchrony, for marking behavioral events.

The software is supported by an expansive online documentation (with tutorial data) and online user forum.

### A tool for augmented research productivity and reproducibility

With IN-Brainstorm, electrophysiologists are provided a free, integrated software environment that promotes and facilitates harmonized principles of data management, methods, documentation, code verification and reproducibility of data analyses. Such practical and user-friendly tools also accelerate the education of electrophysiologist trainees and promotes the adoption and expansion of data harmonization efforts, such as BIDS and *Neurodata Without Borders*.

Every instance of data processing is logged, with the filenames of the data used and time stamps of execution. These simple, yet powerful features document the provenance of data derivatives and analysis results. Custom IN analysis pipelines assembled for elementary processing blocks of the software’s library can be shared with collaborators, publishers and the scientific community. Pipelines are constructed via the GUI and saved as Matlab files. The open-source code of IN-Brainstorm is thoroughly documented, verifiable and can benefit from contributions from any user via GitHub. Sharing is further encouraged and facilitated by Brainstorm’s data organization in Studies, which can be zipped for archiving, exportation (e.g., as a BIDS repository) or importation into the Brainstorm environment of a collaborator. Batch processing of multiple data volumes is automated, thanks to the systematic organization of Brainstorm’s file system and can be executed on high-performance computing servers without requiring GUI interactions.

For all these reasons, we believe that IN-Brainstorm responds to an unmet need of the electrophysiology community. By providing a unique environment with a common set of analytical tools, the application also provides a unique bridge between recording scales, data types and researchers, and additionally, between the methods used in human, animal and slice preparations. It also represents a scalable framework to developments and integration of existing or future tools and data formats for the entire field of electrophysiology.

## Supporting information

Real time LFP activity, time frequency decomposition, and topographical visualization of the recorded activity

## Acknowledgements

We are grateful to Dr. Matthew Krause, Dr. Pedro Vieira, Dr. Christos Gkogkas, Bennet Csorba, Nardin Nakhla and Yavar Korkian for providing datasets. Dr. Shahab Bakhtiari for his input in data analysis. We also thank Elizabeth Bock for early testing of the tools featured. We also extend acknowledgments to Dr. Michael Petrides, Sebastien Tremblay and Veronika Zlatkina for their input.

This work was supported by a Molson Neuro-Engineering Scholarship (the Molson Foundation) to K.N., by grants from the National Science and Engineering Research Council of Canada (NSERC 436355-13 to S.B. and 341534-2012 to C.C.P.), the National Institutes of Health (NIH-1R01EB026299) and the Brain Canada Foundation (PSG15-3755) to S.B.

## Competing Interests

No competing interests declared by the authors.

